# Co-Delivery of Synergistic Antimicrobials with Polyelectrolyte Nanocomplexes to Treat Bacterial Biofilms and Lung Infections

**DOI:** 10.1101/2021.11.22.469570

**Authors:** Joel A. Finbloom, Preethi Raghavan, Michael Kwon, Bhushan N. Kharbikar, Michelle A. Yu, Tejal A. Desai

## Abstract

New approaches are needed to treat bacterial biofilm infections, particularly those of *Pseudomonas aeruginosa* (PA), which have high rates of antimicrobial resistance and are commonly found in chronic wound and cystic fibrosis lung infections. Combination therapeutics that act synergistically can overcome resistance; however, the delivery of multiple therapeutics at relevant dosages remains a challenge. We therefore developed a new nanoscale drug carrier for antimicrobial co-delivery by combining approaches from polyelectrolyte nanocomplex (NC) formation and layer-by-layer electrostatic self-assembly. This strategy led to NC drug carriers loaded with tobramycin antibiotics and antimicrobial silver nanoparticles (AgTob-NCs). AgTob-NCs displayed synergistic enhancements in antimicrobial activity against both planktonic and biofilm PA cultures, with positively charged NCs leading to complete biofilm eradication. NCs were evaluated in mouse models of lung infection, leading to reduced bacterial burden and improved survival outcomes. This approach therefore shows promise for nanoscale therapeutic co-delivery to overcome antimicrobial resistant bacterial infections.

## Introduction

Antimicrobial resistance (AMR) is a global health crisis recognized by the CDC and the WHO as a dire threat to human health. Increasing rates of AMR bacterial infections are outcompeting the development of new antibiotics, and deaths from AMR infections are estimated to increase to over 10,000,000 annual deaths by 2050 (*1*). A major mechanism of antimicrobial resistance in bacterial infections is the development of biofilms, where bacteria encase themselves within heterogeneous biological hydrogels of extracellular DNA, polysaccharides, and protein filaments (*2*). Biofilm formation directly increases bacterial virulence and leads to AMR mechanisms such as limited antibiotic penetration and increased expression of efflux pumps (*2*–*4*). Bacterial biofilms are estimated to occur in up to 80% of human infections and can be 1000 times more resistant to antibiotics when compared to planktonic bacteria (*2*). Biofilm infections are particularly prevalent within chronic wounds such as diabetic skin ulcers, as well as in lung infections of cystic fibrosis (CF) patients. In the lungs of CF patients, the viscous mucus layer caused by ineffective mucus clearance by epithelial cells poses an additional challenging barrier to effective bacterial treatment (*5*). CF patients are thus chronically colonized by bacterial biofilms, particularly those of *Pseudomonas aeruginosa* (PA) bacteria, which once established are nearly impossible to eradicate even in the era of CFTR modulators (*6*–*8*). Chronic colonization by *P. aeruginosa* greatly impacts CF patients’ morbidity and mortality to such extents that continuous month-long courses of inhaled tobramycin and inhaled aztreonam antibiotics are often used to suppress pathologic *P. aeruginosa* strains from developing even when the patient is well. If hospitalized for an exacerbation due to *P. aeruginosa*, CF patients undergo double-coverage with two IV antibiotics of distinct classes for two weeks, making them particularly vulnerable to developing colonization with AMR bacteria.

The co-delivery of synergistic antimicrobial drugs could improve the treatment of AMR bacterial biofilm infections and help eradicate persistent chronic infections (*9*–*11*). These drug combinations can function in parallel to target orthogonal mechanisms and prevent the evolution of resistance pathways. Combination therapeutics can also work together to enhance a singular pathway, such as by facilitating increased uptake of an antibiotic for improved efficacy (*12, 13*). One particularly promising combination of antimicrobial therapeutics is tobramycin and silver nanoparticles (AgNPs). Tobramycin (Tob) is an aminoglycoside antibiotic that is the backbone of most antibiotic therapies used in CF exacerbations due to PA. Several studies have demonstrated that tobramycin and AgNPs act synergistically to overcome resistance of PA biofilms (*14*–*17*). While the exact mechanism of action has not yet been elucidated, it is hypothesized that AgNPs inhibit biofilm formation, increase membrane permeability for enhanced Tob uptake, and generate reactive oxygen species for direct bactericidal activities. Despite its potential, one major challenge to antimicrobial co-delivery is maintaining local therapeutic concentrations of both drugs at the same site of infection, particularly if the site is difficult to reach. This results in immense burdens for patients, who are frequently exposed to prolonged courses of antibiotics at potent doses, which increases patients’ risk for developing adverse reactions and AMR (*5*). In colonized CF patients, inhaled tobramycin and aztreonam are effective in reducing exacerbations due to *P. aeruginosa*, however these therapies require dosing two to three times a day, which encumber patients’ ability to attend work and school. In chronic osteomyelitis, a duration of 2-6 weeks of parenteral antibiotics followed by 4-8 additional weeks of oral antibiotics is required and even then, the infections often persist (*18*).

Therefore, new approaches are needed to enable therapeutic co-delivery, improve antimicrobial efficacies, and overcome resistant bacterial biofilm infections.

Nanoparticle-based strategies for antimicrobial delivery have seen early preclinical success in sustained drug release and treating bacterial infections (*19*–*26*). Polymeric particles are widely used as nanoscale drug carriers for antimicrobial delivery, as they can be fabricated from biocompatible polymers, loaded with bioactive cargo, and delivered via inhalation (*27, 28*). Additionally, polymeric particles have tunable properties such as size, shape, surface charge, and polymer composition, which can influence particle biodistribution, half-life, and interactions with biological barriers such as mucus and bacterial biofilms (*29*–*33*). Polyelectrolyte nanocomplexes (NCs) are an emerging class of polymeric nanoparticle drug carrier, and are formed through electrostatic interactions between oppositely charged components (*34, 35*). These NC drug carriers offer physicochemical tunability dependent on the polymer ratios, chemical structures, pKa’s, and molecular weights (*34*). NCs have further advantages in their ease of fabrication without specialized equipment, and in their high loading efficiencies of therapeutic cargos, especially for drugs of high charge densities (*35, 36*), which often have low encapsulation efficiencies using other polymeric systems (*37*). However, NC formulations for the co-delivery of synergistic antimicrobials remains a challenge, as it is difficult to load diverse classes of cargo within a single NC drug carrier.

To enable the co-delivery of tobramycin and AgNP antimicrobials to treat PA biofilms, we developed a new nanoscale drug carrier fabrication platform that combines approaches from polyelectrolyte nanocomplexation and layer-by-layer (LbL) electrostatic self-assembly. This new NC-LbL strategy led to the formation of NCs co-loaded with Tob and AgNPs (**Figure 1a**). These AgTob-NCs were fabricated from commercially available materials without the use of any specialized equipment. Additionally, AgTob-NCs offer tunability in particle size and surface charge to engineer the drug carrier biointerface and enhance antimicrobial activity. AgTob-NCs had high degrees of loading for both Tob (>75%) and AgNPs (>95%) and facilitated the co-delivery of Tob and AgNP antimicrobials to eradicate PA biofilms that were resistant to either treatment alone. AgTob-NCs demonstrated potency *in vivo* in reducing bacterial burden and improving survival outcomes in mouse models of acute PA lung infection. This approach could see widespread use in enabling the co-delivery of diverse classes of antimicrobials to overcome recalcitrant biofilms such as those present in CF lung infections.

**Fig. 1.**
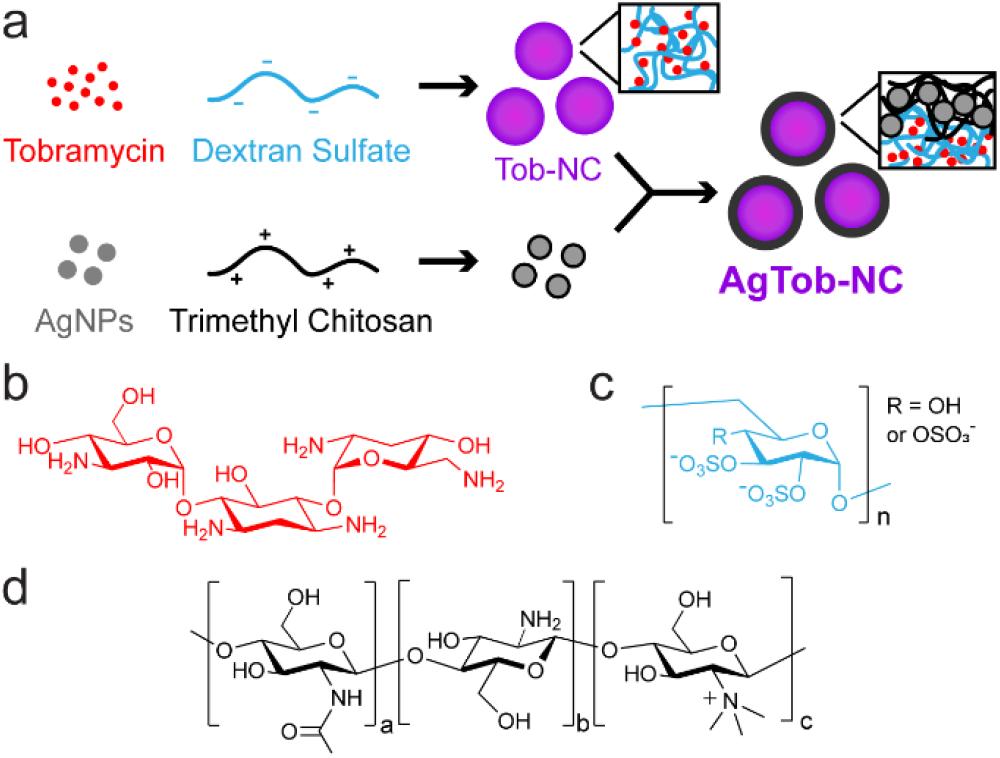
Polyelectrolyte nanocomplex (NC) formation coupled with layer-by-layer (LbL) assembly enables the co-delivery of antimicrobials to treat AMR bacterial biofilm infections. (a) Complexation of tobramycin (positive charge) with dextran sulfate polymers (negative charge) led to the formation of NCs. The LbL addition of trimethyl chitosan and silver nanoparticles (AgNPs) facilitated the co-loading of antimicrobials onto NCs. Chemical structures of (b) tobramycin, (c) dextran sulfate, and (d) trimethyl chitosan (TMC). TMC contains units of chitin, chitosan, and TMC within the polymer, with TMC being the dominant unit.

## Results

### Tobramycin Nanocomplex Fabrication and Characterization

To fabricate nanocomplexes (NCs) co-loaded with tobramycin and silver nanoparticles, NCs were first optimized to encapsulate tobramycin alone, followed by electrostatic layer-by-layer self-assembly with AgNPs (**Figure 1a**). Tobramycin is an aminoglycoside antibiotic with five potential sites for amine protonation (**Figure 1b**) and is therefore well-suited for electrostatic incorporation into polyelectrolyte NCs. Dextran sulfate (DS) was chosen as the complementary negatively charged polymer to form Tob-NCs, as its polysaccharide structure and dense negative charge (**Figure 1c)** make it ideal for complexation with positively charged tobramycin. Tobramycin and dextran sulfate were co-incubated together to form nanocomplexes, which were characterized for drug encapsulation efficiencies as well as particle size and surface charge by dynamic light scattering and zeta potential measurements, respectively (**Figure 2a**).

**Fig. 2.**
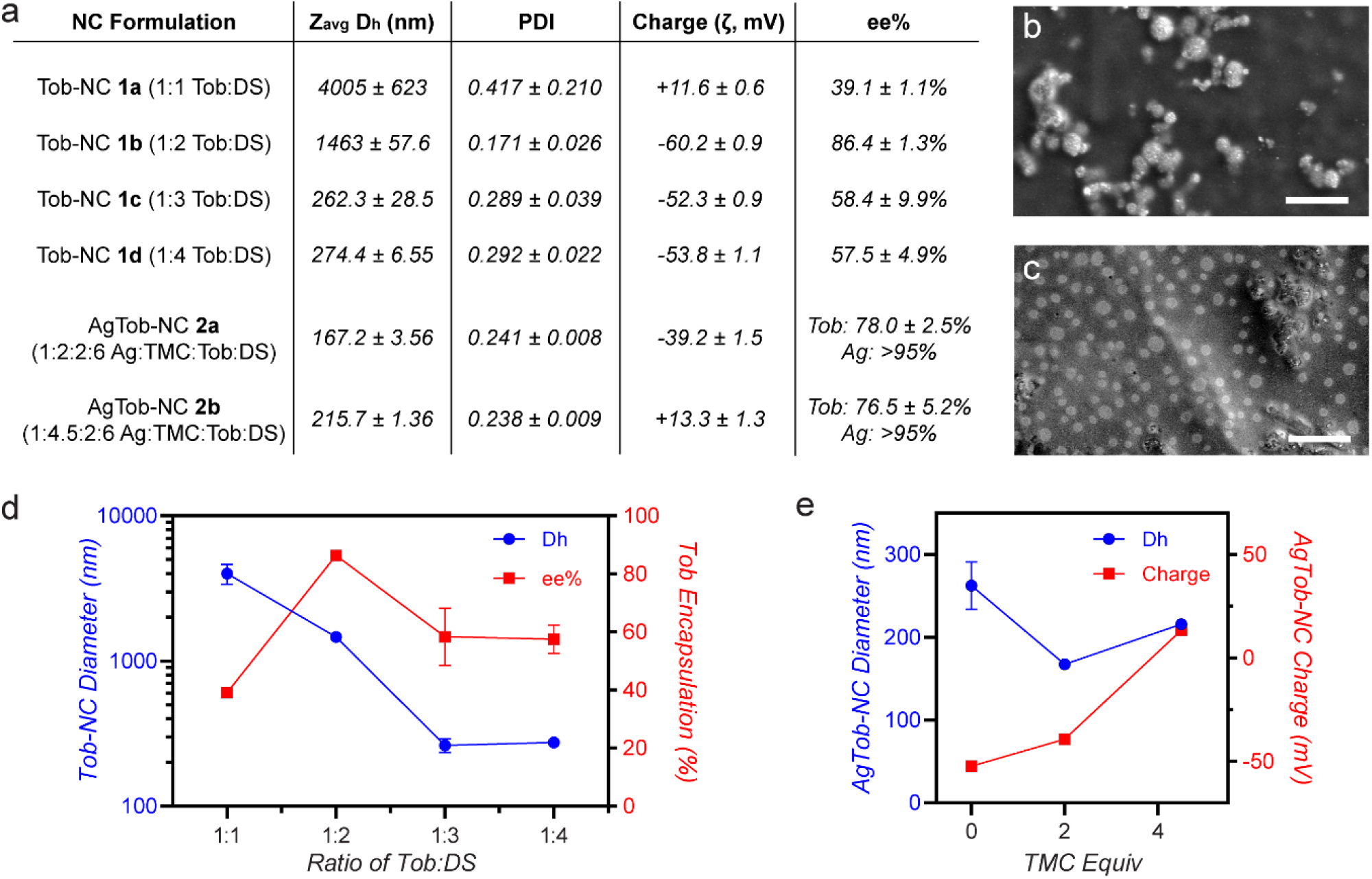
Characterization of antimicrobial-functionalized nanocomplexes (NCs). (a) Hydrodynamic diameter (D_h_, Z_avg_), polydispersity index (PDI), surface charge, and encapsulation efficiency (ee%) measurements for NC formulations, with varying ratios of tobramycin (Tob), dextran sulfate (DS), silver nanoparticles (AgNP), and trimethyl chitosan (TMC). Tob-NC 1c was used to fabricate AgTob-NCs. Both NC **2a** and **2b** demonstrated near-quantitative loading of AgNPs with a 1:2 mass ratio of Ag:Tob. NC 2b was functionalized with an additional 2.5 equiv of TMC to create positively charged particles. (b) SEM image of **2a**. (c) SEM image of **2b**. Scale bars = 2 µm. (d) Tob-NC design parameters with Tob:DS dependent trends for NC size and Tob encapsulation efficiency. (e) AgTob-NC design parameters for TMC-dependent NC size and charge.

NC formation conditions were optimized for pH and buffer type, with optimal conditions observed using 50 mM sodium acetate buffer at pH 4. This pH allowed for increased protonation of tobramycin amines while maintaining DS negative charge. Tob and DS co-incubation at pH 4 demonstrated increased particle formation when compared to pH 6, wherein no Tob encapsulation or particle formation was observed. Using these optimized conditions, we observed Tob:DS ratio-dependent trends in particle size, surface charge, and Tob encapsulation efficiencies (**Figure 2d**). A mass ratio of 1:1 Tob:DS led to microparticles of positive charge that were highly polydisperse and had low Tob encapsulation efficiencies (ee%) of 39%. As the ratio of Tob:DS increased to 1:2, a significant increase in ee% was observed at 86%, however the particles were of ∼1.4 µm in diameter, likely too large to penetrate efficiently through bacterial biofilm hydrogels (*29, 38*). At Tob:DS ratios of 1:3 and 1:4, 250-300 nm particles were formed, with encapsulation efficiencies at 58% (1:3) and 57% (1:4). Thus, Tob-NC **1c**, corresponding to a Tob:DS ratio of 1:3 was chosen for subsequent AgNP loading and antimicrobial studies.

### AgTob-NC Fabrication and Characterization

Tob-NCs **1b**-**1d** displayed strong negative charges (**Figure 2a**), due to the mass ratios of <1 Tob:DS within the particles. As AgNPs also have a negatively charged surface owing to their citrate coatings, we employed a layer-by-layer (LbL) self-assembly approach to co-functionalize NCs with both Tob and AgNPs. LbL approaches have been used extensively in the field of drug delivery and nanoparticle functionalization, although predominantly with thin films and non-NC formulated polymeric particles (*25, 39*–*41*). Commercially available 10 nm AgNPs were chosen for LbL functionalization of Tob-NCs, as we hypothesized that this size would allow for formation of a thin layer on the particle surfaces and have been previously shown to have improved antimicrobial activity when compared to AgNPs of larger size (*14*). Trimethyl chitosan (TMC) was chosen as the positively charged polyelectrolyte to facilitate LbL loading of AgNPs onto nanocomplexes (**Figure 1a**). TMC is a positively charged polysaccharide (**Figure 1d**), which is biocompatible and has mucoadhesive properties, making it particularly attractive for applications in both oral and pulmonary drug delivery (*42*–*44*). Additionally, chitosan and its derivatives have been shown to have antibacterial activities and enhance antibiotic efficacy (*45, 46*). TMC was first incubated with AgNPs at a mass ratio of 1:1 to initiate complexation and form positively charged AgNPs. TMC-coated AgNPs were then added to Tob-NCs to form AgTob-NCs and additional TMC was added to stabilize the particles and prevent aggregation. Using this method at Na acetate pH 4, quantitative loading of AgNPs onto NCs was observed, with tunable mass ratios of Ag:Tob ranging from 1:2 to 1:8. A ratio of 1:2 Ag:Tob was chosen to maximize loading and synergistic antimicrobial activities. Interestingly, the immediate addition of AgNPs to freshly formed Tob-NCs led to a 50-100 nm decrease in particle diameter (**Figure 2a**,**e**). This phenomenon is likely due to the strong electrostatic attraction between TMC and DS, as has been observed previously (*44*). This strong complexation also led to an increase in overall Tob ee% for AgTob-NCs, increasing from 58% for Tob-NC **1c** to >75% encapsulation efficiencies for all AgTob-NCs fabricated.

In addition to co-loading AgNPs and Tob onto a singular nanocarrier, we were interested in modulating the physicochemical properties of these NCs to design their biointerfacial interactions with bacterial biofilms. As *P. aeruginosa* biofilms are composed of negatively charged biopolymers such as DNA and alginate, we hypothesized that positively charged NCs of a 200-300 nm size regime could initiate interactive filtering within biofilms (*29, 38*), with their size enabling steric diffusion through biofilm hydrogel pores while still allowing for electrostatic engagement with the biofilm matrix to increase NC attachment and antimicrobial delivery. To create positively charged AgTob-NCs, additional equivalents of TMC were added following AgNP co-incubation. These studies yielded two different NC formulations, AgTob-NC **2a** bearing a negatively charged surface, and AgTob-NC **2b** with a positively charged surface (**Figure 2a**,**e**). Both NCs **2a** and **2b** had similar degrees of Tob ee%, with **2b** being of slight increased size, likely owing to the additional TMC layers coating its surface.

### Storage, Stability, and Biocompatibility of AgTob-NCs

AgTob-NCs **2a** and **2b** were evaluated for their long-term storage potential, stability in biological environments, and biocompatibility. Both NC **2a** and **2b** were able to be lyophilized and resuspended without significant aggregation or destabilization, although a slight increase in particle size to ∼300 nm in diameter was observed upon resuspension (**Figure S2**). To determine whether the NCs would destabilize or aggregate in biological media, NCs were incubated in 10% FBS for 20 h at 37 °C prior to characterization via DLS. NC **2a** maintained its size and no aggregation was observed, while **2b** increased in size to ∼1 µm (**Figure S2**), likely owing to modest aggregation of NCs through the adsorption of serum proteins onto the positively charged particle surfaces. Presto Blue cell viability studies were performed using A549 lung cells incubated with AgNPs or NCs for 24 h. AgNPs can be cytotoxic at higher concentrations (*47, 48*), however we observed only modest decreases in cell viability to ∼85% with AgNPs or NCs **2a** and **2b** at the highest dosage tested of 4 µg AgNP per 20,000 cells, with no significant differences measured between groups (**Figure S3**). Thus, AgTob-NCs demonstrated favorable properties for long-term storage, stability, and biocompatibility.

### Antimicrobial Activity of AgTob-NCs Against Planktonic *P. aeruginosa*

With both NC formulations in hand, we set out to evaluate whether NC-mediated co-delivery of Ag and Tob provided improved antimicrobial activity against planktonic cultures of *P. aeruginosa*. Laboratory strain PA14 was used for all antimicrobial activity assays, as PA14 is a well-studied virulent PA strain and can be cultured to form robust biofilms (*49, 50*). Tobramycin, AgNPs, and AgTob-NC **2a** and **2b** were cultured with dilute concentrations of PA for 20 h to determine the minimum inhibitory concentration (MIC) of tobramycin required to inhibit 80% of planktonic bacterial growth for each group relative to untreated PA14 cultures (**Figure 3**).

**Fig. 3.**
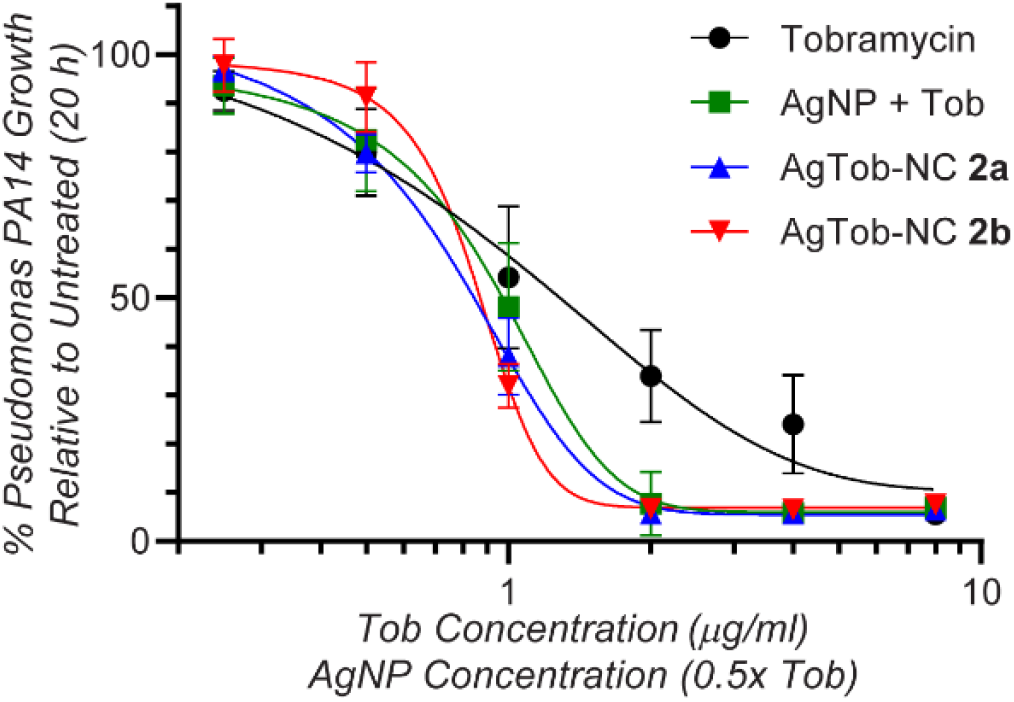
Antimicrobial activity of AgTob-NCs against planktonic *Pseudomonas aeruginosa* cultures. PA14 laboratory strains were incubated with Tob, AgNPs+Tob, or AgTob-NC formulations for 20 h while shaking, prior to measuring bacterial growth at OD600. The minimum inhibitory concentration (MIC80) of tobramycin decreased from 8 µg/ml to 2 µg/ml with AgTob-NC treatment, demonstrating effective co-delivery and synergistic enhancement in antimicrobial activity.

Tobramycin alone had an MIC value of 8 µg/ml, which is in agreement with the range commonly reported in the literature (*13, 51*). When coupled with AgNPs either co-incubated in solution (AgNP + Tob control) or co-loaded into NCs **2a** and **2b**, a significant enhancement in antimicrobial activity was observed, with Tob MIC values decreasing to 2 µg/ml (**Figure 3**). AgNPs alone had an MIC value of 4 µg/ml (data not shown), leading to a fractional inhibitory concentration (FIC) index value of 0.5, confirming synergistic activity as defined by FIC ≤ 0.5 (*52*). No significant differences were observed between any of the Ag+Tob treatment groups. Thus, NCs **2a** and **2b** delivered both antimicrobials effectively without sacrificing drug potency and enhanced the inhibition of planktonic *P. aeruginosa* growth when compared to Tob or AgNP treatments alone.

### Antimicrobial Activities of AgTob-NCs Against *P. aeruginosa* Biofilms

As the majority of PA infections exist as biofilms that are associated with high rates of AMR, we next evaluated the antimicrobial activities of AgTob-NCs against PA14 biofilms. Biofilms were cultured for 24 h using previously reported protocols prior to the addition of therapeutics and further incubation for 20 h (*50*). PA14 biofilms were stained using a BacLight live/dead bacterial stain, with Syto9 (S9) acting as a universal stain for all bacteria and propidium iodide (PI) as a stain for dead bacteria with permeable membranes. PA14 grew into robust biofilms that displayed significant antimicrobial resistance to tobramycin, even at higher dosages of 40 µg/ml, corresponding to 6.4 µg of Tob per well (**Figure 4a**). AgNP treatment alone also demonstrated no observable antimicrobial or antibiofilm activities (**Figure S4**).

**Fig. 4.**
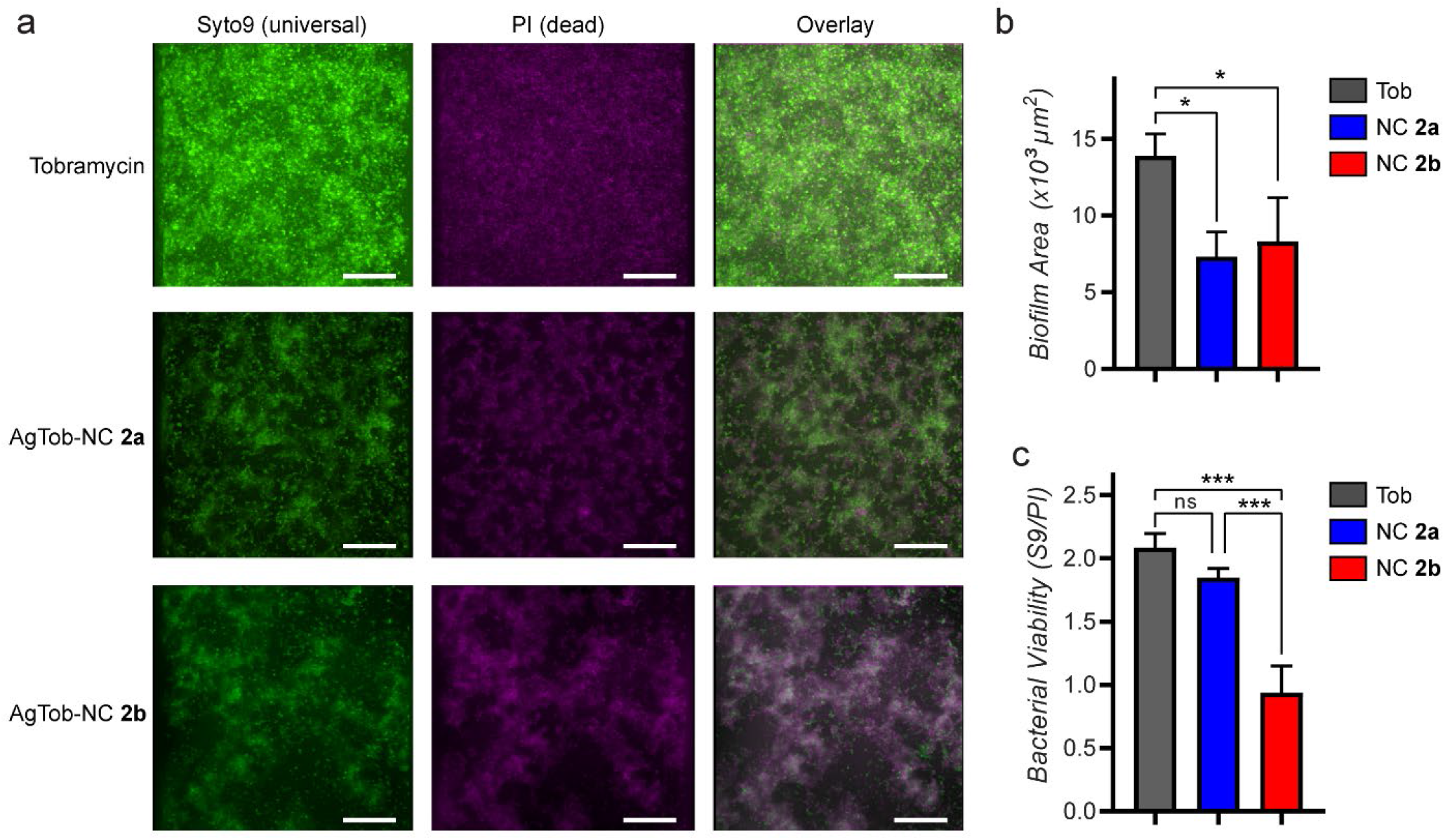
Antibiofilm activities of AgTob-NCs. (a) Representative 3D projections of confocal fluorescence microscopy images of PA14 biofilms treated with Tob, AgTob-NC **2a**, or AgTob-NC **2b**. All biofilms were treated with 6.4 µg of Tob and/or 3.2 µg AgNP for 20 h. Syto9 (green) is a universal stain for both live and dead bacteria, while PI (magenta) stains only dead bacteria. White indicates overlay at approximately equal fluorescence intensities. Tobramycin alone did not demonstrate any significant antibiofilm activity with most dead bacteria observed outside of biofilm regions, while NC-mediated co-delivery of AgNPs and Tob was able to disrupt biofilm structures, with NC **2b** demonstrating the most pronounced antibiofilm and bactericidal activities. Scale bars = 25 µm. Images obtained at 40x magnification. (b,c) Quantification of biofilm images to determine (b) biofilm biomass areas and (c)bacterial viability as measured through the ratio of S9/PI intensities within biofilms. Image quantification was performed using 20x magnification images. * p<0.05, *** p<0.001. Additional images of biofilms treated with all groups and controls at 20x magnification are available in the Supporting Information.

While neither tobramycin nor AgNP treatments alone could reduce biofilm formation or cause significant reductions in bacterial viability, both AgTob-NCs **2a** and **2b** treatments led to observable disruptions in biofilm morphologies and significant reductions in bacterial biomass (**Figure 4b**). Images were quantified to determine the ratios of S9/PI fluorescence intensity as a marker for bacterial viability within biofilms. Only NC **2b** led to a significant loss in bacterial viability within biofilms, with S9/PI ratios decreasing to ∼1.0, indicating that all the bacteria present within the biofilms were dead (**Figure 4c**).

As the predominant difference between NC formulations was in the increased TMC coating and positive charge of NC **2b**, we sought to elucidate the influence of surface charge on the distribution of NCs within PA biofilms, as we hypothesized that the positively charged NC **2b** would have increased interactions with the negatively charged biofilm extracellular matrix. Fluorescently labeled NCs **2a** and **2b** were fabricated using fluorescein-conjugated dextran sulfate (FITC-DS) and were added onto PA14 biofilms for 1 h prior to analysis of NC-biofilm interactions via confocal fluorescence microscopy (**Figure 5**). Microscopy analysis revealed that NC **2b** occupied nearly twice the area within biofilms when compared to NC **2a** (**Figure 5c**), and this increase in biofilm distribution likely led to enhanced antimicrobial activity.

**Fig. 5.**
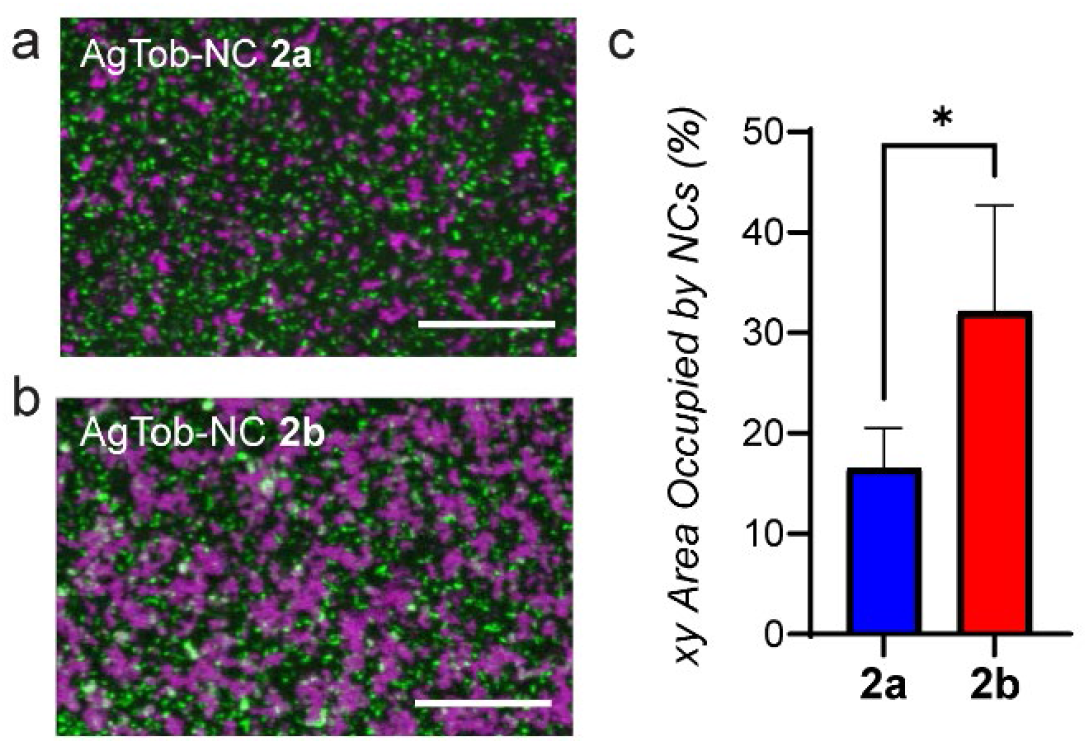
Distribution of fluorescently labeled AgTob-NCs within PA14 biofilms. (a,b) Representative 3D projections of confocal fluorescence microscopy images of FITC-NCs (magenta) within biofilms (green). Images were acquired 1 h after NC addition to biofilms. Scale bars = 50 µm. (c) Quantification of biofilm images to determine % xy area occupied by NCs within biofilms at maximal fluorescence intensity z positions. * p<0.05

### Antimicrobial Activity of AgTob-NCs in Mouse Models of Lung Infection

After determining the efficacy of AgTob-NCs in inhibiting bacterial growth and disrupting biofilms *in vitro*, we next tested the antimicrobial activity of NC **2b** drug carriers in mouse models of CF lung infection. Wildtype C57BL/6 mice were anesthetized under isofluorane for 10 min, then intratracheally instilled with PBS, Tobramycin, or AgTob-NC **2b**. After 2 h, the mice were anesthetized again with isofluorane and challenged via intratracheal instillation with 2×10^6^ colony forming units (CFU) of PA14 per animal and monitored for survival outcomes over 24 h. This equilibration interval was chosen to model inhaled antibiotic therapies in humans, which precede active exacerbations and are used as maintenance therapies, as well as to allow the mice to recover from anesthesia. Physiologic measurements of temperature and weight change were taken every 2 h after instillation, and no mice needed to be excluded based on the IACUC protocol for post-procedural weight loss. Treatment with PBS control prior to infection led to median survival of 8 h, with 0/9 mice surviving the course of the study (**Figure 6a**). When treated with tobramycin, survival outcomes increased to 3/8, while NC **2b** led to the highest survival rates of 8/10 mice (p = 0.0004). Temperature analysis of mice revealed body temperature decreases of approximately 12 °C for the PBS-treated group, whereas Tob and NC treatments led to temperature decreases of approximately 5 °C over the course of the study (**Figure S5**).

**Fig. 6.**
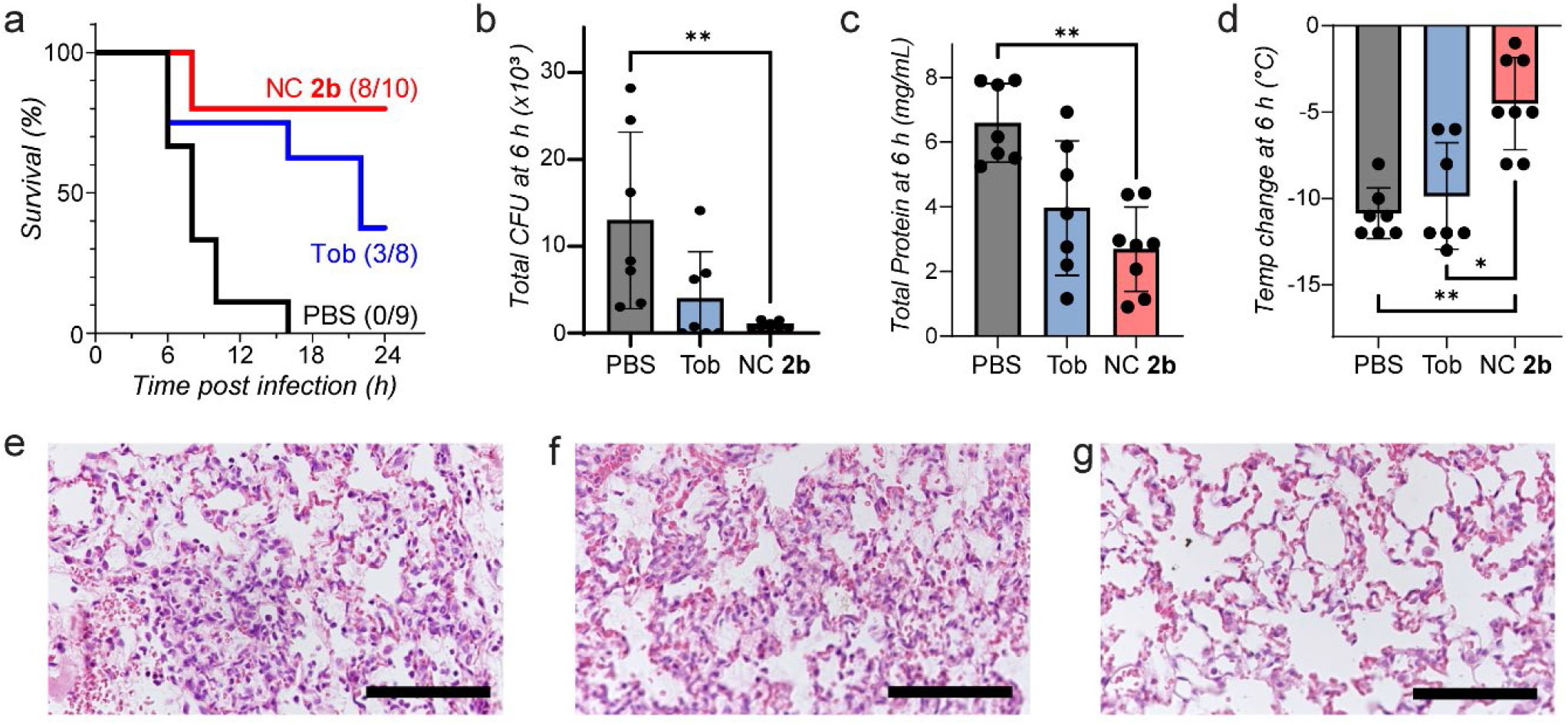
Antimicrobial activity of AgTob-NCs in mouse models of lung infection. Wildtype mice were treated with 0.2 mg/kg tobramycin, AgTob-NC **2b**, or control (PBS) for 2 h prior to challenge with 2×106 colony forming units (CFU) of PA14. Mice were monitored for weight and temperature change, and were either evaluated for survival over 24 h or sacrificed at 6 h post-challenge to quantify bacterial burden and physiological markers of infection. (a) Kaplan Meier curve revealed improved survival outcomes in NC-treated mice, with 8/10 mouse survival compared to 3/8 for Tob and 0/9 for PBS treated groups, respectively (p = 0.0004). (b) Analysis of bacterial burden at 6 h revealed that AgTob-NCs effectively reduced bacterial infection by approximately 1.5 log-fold when compared to PBS control. (c) A reduction in total protein count was determined in both Tob and NC-treated mice. (d)Mice treated with NCs had reduced temperature decreases over 6 h when compared to PBS or Tob treatments. (e-g) Representative images of haematoxylin and eosin (H&E)-stained lung sections for (e) PBS, (f) Tob, and (g) NC-treated mice. Scale bars = 100 µm. For all graphs, * p<0.05, ** p<0.01, *** p<0.001.

To quantify the bacterial burden of mice during the early stage of infection, mice were sacrificed at 6 h post-infection and CFU and physiological markers of infection were assessed. Total CFU counts combined from both bronchoalveolar lavage (BAL) and lung homogenization revealed that NC treatment reduced bacterial burden by ∼1.5 log-fold to levels of near-clearance when compared to PBS control (**Figure 6b**, p = 0.0026). Although no statistically significant differences were observed for white blood cell count or weight gain between groups (**Figure S6**), NC treatment did reduce total protein levels compared to PBS control (**Figure 6c**, p = 0.0024), while also limiting body temperature decreases over 6 h (**Figure 6d**, p = 0.0063). Histological analysis of lung sections after 6 h sacrifice revealed decreased neutrophilic inflammation and lung injury as measured by interstitial edema and disruption of the alveolar epithelial barrier in NC-treated mice (**Figure 6e-f**). For all quantitative metrics of infection, no statistical differences were observed between Tob and PBS treatment groups. Taken together, these studies demonstrate that NC therapy was the most effective treatment at reducing bacterial burden and physiological markers of infection and inflammation and increased overall survival outcomes.

## Discussion

Rising rates of antimicrobial resistant bacterial infections coupled with limited approval of new antibiotics have led to a global crisis and fears of entering a post-antibiotic era. It is therefore imperative to develop new drug carrier formulations capable of enhancing the activities of already-approved antibiotics such as tobramycin. By combining polyelectrolyte nanocomplexation with layer-by-layer electrostatic self-assembly, we developed a new class of nanoscale drug carrier, capable of co-delivering tobramycin antibiotics with antimicrobial silver nanoparticles. While this drug combination had been previously reported in the literature for enhanced antimicrobial activity against PA biofilms (*14*), to our knowledge it had never been tested in a nanoscale drug delivery formulation nor in an *in vivo* model of infection.

Effective dual delivery of antimicrobials is particularly important for the eradication of bacterial biofilms, as insufficient antimicrobial delivery at sublethal doses can in fact enhance biofilm growth (*53, 54*). Indeed, we observed that Tob and AgNP individual treatments led to greater biofilm formation when compared to untreated biofilm controls (**Figure S4**). Although dead bacteria were observed with Tob treatment, PI-stained bacteria were predominantly found outside of biofilm morphologies, indicating that while Tob may be effective against planktonic PA14, it was unable to permeate throughout the biofilm hydrogels to achieve effective antimicrobial activity, as has been previously observed in literature reports (*4, 29*). This further highlights the need to develop nanocarrier formulations capable of interfacing with bacterial biofilms to improve the biofilm permeation and efficacy of antimicrobials.

The drug delivery method developed herein led to high encapsulation rates of both tobramycin and AgNPs into the NC formulation while simultaneously allowing for physicochemical engineering of particle size and surface charge. These parameters are critical for the navigation of particles through biological barriers such as bacterial biofilms (*29*). NCs **2a** and **2b** were developed to match the reported pore sizes of biofilms. As bacterial biofilms are composed of negatively charged biopolymers such as polysaccharides and DNA (*2*), we hypothesized that positively charged NC **2b** would have strong electrostatic interactions with the biofilm matrix, enhancing the engagement and permeation of NC drug carriers throughout the biofilm via interactive filtering. In comparison, we hypothesized that negatively charged NC **2a** would quickly penetrate through the biofilm matrix or only locate to larger water channels within the biofilm hydrogel owing to the electrostatic repulsion between NC **2a** and the negatively charged biofilm matrix. Indeed, when observing NC distribution within established PA biofilms, we observed that NC **2b** occupied nearly double the area as **2a** after 1 h of incubation (**Figure 5**). This increase in distribution likely contributed to the enhanced antibiofilm and bactericidal activity of NC **2b** when compared to **2a** (**Figure 4**). While both NCs could release their payloads as determined in the planktonic MIC assays **(Figure 3**), only NC **2b** readily permeated the PA biofilm to deliver both Ag and Tob throughout the biomass, leading to complete biofilm eradication. These studies underscore the importance of drug carrier biointerface design (*29*), as these considerations can have significant effects on therapeutic outcomes.

Following *in vitro* studies, we set out to evaluate the therapeutic potential of NCs to clear PA lung infections *in vivo*. AgTob-NCs were evaluated in lung infection models as PA infections are prevalent in the lungs of cystic fibrosis patients, where they are notoriously recalcitrant, requiring prolonged antibiotic treatment at high doses, which even then oftentimes does not prevent reinfection (*7*). As NC **2b** demonstrated improved antimicrobial and antibiofilm activities when compared to **2a** in our biofilm studies, it was selected for therapeutic evaluation in mouse models of lung infection. In addition to biofilm permeation properties, the additional coatings of TMC on NC **2b** surfaces could allow for increased bioadhesion and local retention, as TMC and other chitosan derivatives have been shown to have broad cyto- and mucoadhesive properties (*35*). For the model developed for these studies, therapeutics were administered intratracheally 2 h prior to infection. When compared to other commonly used mouse models of lung infection where therapeutics and bacteria are co-administered simultaneously or in quick succession (*20, 21, 55*), this approach tests the sustained effects of therapeutics following administration.

Using this model of lung infection, we observed that NC **2b** reduced bacterial burdens and improve survival outcomes to 80% when compared to PBS and Tob treatments, which had 0% and 37.5% survival outcomes, respectively (**Figure 6**). As our model led to a median survival of 8 h without antibiotic intervention, mice were sacrificed at 6 h to quantify bacterial burden at a relatively early time point within the spread of infection. This provided us with a holistic viewpoint of both the overall outcomes of treatment using the survival study, and insight into the antimicrobial activities and biocompatibilities of Tob and NC **2b** using our 6 h sacrifice model. While tobramycin treatment did not differ significantly from PBS in terms of reducing bacterial burden at 6 h, NC **2b** treatment led to near complete clearance of bacterial load within the lungs. Some Tob-treated mice did display low bacterial load, however the wider distribution of bacterial load within the Tob treatment group suggests that Tob treatment effects were heterogeneous when compared to NC delivery. The NC-improved bacterial reduction likely contributed to the 80% survival outcomes of NC treatment. Furthermore, NC **2b** treatment led to a reduction in lung protein levels at 6 h, indicating reduced lung injury. This finding agrees with histology images, wherein NC-treated lungs had reduced inflammation and healthier alveolar structures when compared to PBS and Tob treated lungs. Although no weight changes were observed between groups, this is likely due to the shorter 6 h sacrifice timepoint, which would limit observable weight differences from the start of the study. Lastly, NC **2b** led to the greatest stability in body temperature when compared to PBS and Tob treatments, indicating that the NC treatment was well tolerated and effective in clearing bacterial lung infections.

These studies were limited to Tob treatment of *P. aeruginosa* as PA is one of the most dominant bacterial species in CF lung infections and Tob is commonly prescribed for PA infections. However, the NC-LbL strategies developed herein are highly versatile and could be applied toward other charged antibiotics such as gentamicin and polymyxins for use against both PA and other bacterial infections in lungs, wounds, and fractures. While the NCs in this study were administered via intratracheal instillation, their size, stability profiles, and ability to be lyophilized could enable their delivery via inhalation, as has been demonstrated for other nanoparticle formulations (*28*). This would increase the translational potential of AgTob-NCs and lead to their incorporation into an inhaler device similar to the commercially available Tobi Podhaler.

In summary, the NC-LbL antimicrobial formulation strategy developed in these studies led to the fabrication of tunable nanocomplexes and facilitated the co-delivery of Tob and Ag to eradicate *P. aeruginosa* in both planktonic and resistant biofilm states, and improved therapeutic outcomes in mouse models of PA lung infection. By engineering the physicochemical properties of the nanocomplexes, we observed how surface chemistry influenced particle biointerfaces and resultant antimicrobial activities. Overall, this approach shows promise in the co-delivery of diverse classes of antimicrobials to treat bacterial infections, including the resistant biofilms associated with cystic fibrosis lung infections.

## Materials and Methods

### General Methods and Instrumentation

Unless otherwise noted, all reagents were purchased from commercial sources. Low molecular weight trimethyl chitosan was purchased from Sigma Aldrich (St. Louis, MO) with a reported >70% degree of quaternization. 40 kDa Dextran sulfate was purchased from Sigma Aldrich (St. Louis, MO). All reagent solutions were freshly prepared for each experiment and fabrication process. Silver nanoparticles of 10 nm diameter were purchased from nanoComposix (San Diego, CA). Glycerol stocks of *P. aeruginosa* PA14 were generously provided by Dr. Oren Rosenberg. A549 lung cells were provided by the UCSF Cell and Genome Engineering Core.

Zeta potential and dynamic light scattering measurements were conducted on a Malvern Zetasizer Nano ZS. Absorbance and fluorescence quantification measurements were conducted on a Molecular Devices SpectraMax M5 plate reader. All fluorescence microscopy studies were conducted at the UCSF Nikon Imaging Center using a Nikon Ti spinning disk confocal microscope. Scanning Electron Microscopy (SEM) images were obtained at the UCSF Bioengineering & Biomaterials Correlative Imaging Core using a Zeiss Sigma 500 VP (Carl Zeiss Microscopy GmbH).

### Tob-NC and AgTob-NC Fabrication

Tobramycin and dextran sulfate (DS) were prepared as 20 and 40 mg/mL solutions in ddH_2_O, respectively. NC formation was accomplished through stepwise addition of DS followed by Tob into a solution of 50 mM Na acetate pH 4 for a final concentration of 2 mg/mL Tob and variable concentrations of DS to achieve mass ratios ranging from 1:1 to 1:4 Tob:DS. The solution was mixed by pipetting and left to incubate for at least 5 min before any subsequent reactions or characterizations.

For all AgTob-NC formulations, Tob-NCs were first fabricated at a 1:3 Tob:DS mass ratio. AgTob-NCs were typically fabricated at a 300 µL scale. Prior to fabrication, TMC (7.5 µL of 4 mg/mL in Na acetate pH 4) and AgNPs (30 µL of 1 mg/mL stock solution obtained from commercial sources) were incubated together to achieve a 1:1 mass ratio. Next, a 75 µL solution of 0.05% (w/v) tween was prepared in ddH_2_O, to which were added Tob-NCs (30 µL, 2 mg/mL), followed by the TMC-AgNP solution (37.5 µL). Additional TMC was then added as a 45 µL solution in ddH_2_O (1 equiv for NC **2a**, 3.5 equiv for NC **2b**), followed by 60 µL of 5% (v/v) of poly(vinyl alcohol) and 52.5 µL ddH_2_O to obtain AgTob-NCs as a 300 µL solution at Tob and Ag concentrations of 200 µg/mL and 100 µg/mL, respectively. Tween was used to prevent NC aggregation, while PVA was incorporated to enable the lyophilization of NCs for long-term storage. AgTob-NCs were either used immediately or flash-frozen and lyophilized.

### NC Characterization and Drug Loading Determination

Dynamic light scattering (DLS) and zeta potential measurements were conducted using a Malvern Zetasizer Nano ZS. NCs were typically analyzed at a Tob concentration of 50-100 µg/mL. Particle hydrodynamic diameters (D_h_) were reported using Z_avg_ measurements. Full DLS curves were obtained for intensity size distributions (**Figure S2**). Encapsulation efficiencies (ee%) for Tob were determined by analyzing the NC supernatants following centrifugation (13,000 rpm for 5 min) to evaluate drug depletion after NC formation and LbL AgNP loading. Centrifugation at this speed and time allowed for the formation of an NC pellet without pelleting AgNPs or Tob. AgNP depletion was analyzed via UV-Vis absorbance at 390 nm, while Tob depletion was assessed via an *o*-phthalaldehyde fluorescence assay (*37*). Briefly, *o-*phathalaldehyde was prepared as a 1 mg/mL solution in 100 mM Na borate pH 10.4 buffer with the addition of MeOH for 10% (v/v) and β-mercaptoethanol for 0.5% (v/v) final. *o-*Phthalaldehyde solutions (15 µL) were added to iPOH (90 µL) prior to the addition of sample supernatant (15 µL). Solutions were incubated for 20 min and then analyzed for fluorescence intensity (360/450 ex/em) and compared to Tob calibration curves. *o-*Phthalaldehyde stock solutions could be stored at 4 °C for 1-2 weeks without any observable loss in signal.

The morphology of the NC suspensions was determined using field emission scanning electron microscope (FE SEM), Zeiss Sigma 500 VP (Carl Zeiss Microscopy GmbH). Samples for electron microscopy were prepared by mounting droplets of NC suspensions (20 µL) onto carbon substrate-coated SEM stubs and allowed to air dry. Subsequently, samples were stained with Uranyless. The Uranyless droplet was placed on the hydrophobic surface (parafilm). The stubs with NCs’ side were placed on the Uranyless drop for 2 minutes and thereafter blotted by filter paper to remove access stain on the NCs. Followed by washing 3 times with ddH_2_O at room temperature. Finally, the samples were dried at room temperature for at least 12 h prior to analyze via FE SEM.

### Mammalian Cell Culture and Biocompatibility Studies

A549 cells were cultured in F12 growth medium containing L-glutamine, supplemented with 10% fetal bovine serum and penicillin/streptomycin, and incubated at 37 °C and 5% CO_2_. For biocompatibility studies, A549 cells were trypsinized, resuspended to a concentration of 100,000 cells/mL, and plated in triplicate into a 96 well plate for a final cell number of 10,000 cells/well. After 24 h incubation, the media was removed and replaced with 200 µL of media + treatment (AgNP, AgTob-NC **2a**, or AgTob-NC **2b**). Treatments were prepared by fabricating the nanocomplexes as previously described at concentrations of 200 µg/mL Tob and 100 µg/mL Ag in a final volume of 300 µL. This was further diluted into A549 media for a final Ag concentration of 20 µg/mL, the highest concentration experimental condition. This was serially diluted 1:2 in A549 media until reaching 0.625 µg Ag/mL and added to the cells. The cells were then incubated for an additional 24 h. Following incubation, 20 µL of Presto Blue were added to each well. The cells were then incubated for 1 h and Presto Blue fluorescence intensity (ex/em 560/590) was measured to determine cell viabilities relative to untreated cells.

### Planktonic *P. aeruginosa* Antimicrobial Activity Studies

PA14 was cultured overnight from frozen glycerol stocks at 37°C and 220 rpm in LB media. PA14 cultures were then refreshed by performing a 1:8 dilution into LB and shaken for an additional 1-3 h until an OD_600_ > 0.4 was achieved to confirm log-phase bacterial growth. PA14 was then plated into 96-well plates at 100 µL and OD_600_ = 0.002 in accordance with previously reported methods (*56*). Next, 100 µL of treatment groups and controls were added per well at 2x concentration to achieve PA14 OD_600_ = 0.001 and treatment concentrations starting at 8 µg/mL Tob and 4 µg/mL Ag with 1:2 serial dilutions to a final Tob concentration of 0.25 µg/mL. All antimicrobial studies were performed with a minimum of n=3 biological replicates. Plates were then incubated at 37°C and 220 rpm for 20 h prior to measurement of OD_600_ to assess bacterial viability relative to untreated controls. MIC values are defined as the minimum concentration of drug required to inhibit 80% bacterial growth relative to untreated controls.

### *P. aeruginosa* Biofilm Studies

PA14 biofilms were grown using previously reported methods (*50*). PA14 overnight cultures were refreshed by performing a 1:8 dilution into LB and shaken for an additional 1-3 h until an OD_600_ > 0.4 was achieved to confirm log-phase bacterial growth. Bacteria were then plated into half-area well plates (Greiner Bio-One) at 100 µL and OD_600_ = 0.02 and sealed with air-permeable “Breathe-Easy” membranes. Plates were then placed into a stationary incubator with water bath and incubated at 37 °C for 24 h. After 24 h, the air-permeable membranes were removed, and treatment groups were added to biofilms at 4x concentrations and 40 µL per well. Next, 20 µL of BacLight dyes Syto9 and propidium iodide (co-incubated at 1:300 dilutions each from purchased stocks into PBS following manufacturer’s instructions) were added to each well. Care was taken to add solutions dropwise to the top of each well without disturbing the biofilm. After treatment and BacLight dye addition, plates were sealed with air-permeable membranes and incubated at 37 °C for 20 h with gentle rotations at 50 rpm. After 20 h, bacterial biofilms were imaged using confocal fluorescence microscopy, with S9 excited using a FITC channel and PI excited using a Cy3 channel. Images were acquired using both 20x and 40x air objectives. Images were acquired as z-stacks of biofilms, with 2 µm spacing and 50 µm total slices. Z-stacks were centered at the z-position of highest S9 fluorescence intensity and relative z-stacks were obtained ± 25 µm from that slice. All biofilm studies were performed with a minimum of n=3 biological replicates.

### Biofilm Image Analysis and Quantification

Image quantification of biofilm area and viability was done through ImageJ. First, the background was subtracted from all images within a stack, based upon the z-slice that had the brightest S9 signal. A universal threshold value was applied to all images within a z-stack in order to segment biofilm mass from planktonic bacteria. This threshold was calculated based on Otsu’s binarization of the S9 channel in the untreated control biofilm. Within each segmented area of biomass, the average fluorescence intensities of S9 and PI were measured. This process was repeated for the 5 z-slices above and below the brightest S9 z-slice in a stack. Biofilm area, S9 fluorescence intensity, and PI fluorescence intensity were averaged from these 11 slices for each biofilm z-stack. Image analysis was conducted on z-stacks from 3 biological replicates and plotted as mean ± standard deviation. Statistical analysis was performed using one-way ANOVA with multiple comparisons (Tukey’s method).

### Care and Use of Mice

Mice were housed and bred in specific pathogen-free housing at the UCSF Laboratory Animal Research Center. All experiments were in accordance with the ethical principles and guidelines approved by the UCSF Institutional Animal Care and Use Committee (IACUC). 8–12-week-old male C57BL/6J wildtype mice were obtained from the Jackson Laboratory and used for lung infection studies. Pilot experiments with both female and male WT mice showed no difference between genders, hence male mice were selected for this experiment.

### Intratracheal PA Models of Lung Infection

Mouse models of lung infection were performed according to established techniques (*49*). PA14 was streaked on plates from frozen glycerol stocks and a single colony selected for overnight culture at 37°C and 220 rpm in TSB media. PA14 cultures were then refreshed the morning of infection by performing a 1:8 dilution into TSB and shaken for an additional 3 h until an OD_600_ > 0.4. Mice were randomly distributed into three groups (PBS, Tob, NC treatments) and anesthetized with isofluorane prior to intratracheal instillation of therapeutics. Treatment groups were delivered at dosages of 0.2 mg/kg Tob and 0.1 mg/kg Ag in a total of 50 μL injection volume. Mice were allowed to recover for 2 h prior to isofluorane anesthetization and intratracheal instillation of 2 × 10^6^ CFU PA14 per animal in a volume of 50 μL. Animal weights and rectal temperatures were monitored every 2 h for survival studies or at 6 h prior to euthanasia for quantitative bacterial load and pathophysiology determination. To evaluate bacterial load at 6 h, a bronchoalveolar lavage (BAL) was performed prior to sacrifice and lung collection. BAL samples were used to determine BAL bacterial CFU, white blood cell (WBC) numbers, and total protein levels. Lungs were homogenized in 1 mL of sterile PBS using sterile 100 µm filters and the sterile plunger from a 3 mL syringe. 50 uL of lung homogenate was then serially diluted and plated on PIA plates. CFU was calculated from dilution plates with 30-300 colonies, dividing the colony number by the dilution factor.

Total CFU was calculated by adding BAL and lung homogenate CFU counts. In the cases in which multiple dilution plates had 30-300 colonies, the average between plate counts was used. White blood cells (WBCs) were measured by a hematology analyzer (Genesis; Oxford Science). Statistical analysis of survival studies was performed using a Mantel-Cox long rank test. Statistical analyses of quantitative metrics of bacterial infection and pathophysiology (CFU counts, WBC, total protein, weight change, temperature change) were performed using Kruskal-Wallis nonparametric analyses with Dunn’s correction for multiple comparisons. All statistics were performed using GraphPad Prism 9.

## Author contributions

Conceptualization: JAF, MAY, TAD

Material development and characterization: JAF, PR, BK

Evaluation of antimicrobial and antibiofilm activity in vitro: JAF, PR

Evaluation of antimicrobial activity in vivo: JAF, MK, MAY

Writing—original draft: JAF

Writing—review & editing: JAF, PR, MAY, TAD

## Data and materials availability

Additional data that support the findings of this study are available from the corresponding authors upon reasonable request.

## Supplementary Materials

Figures S1-S6. SEM of Tob-NCs, AgTob-NC stability studies, Biocompatibility studies, Biofilm images at 20x magnification, *In vivo* bacterial load and physiological markers for survival study and at 6 h sacrifice

