## Supplemental Figures S1-S6 for "Co-Delivery of Synergistic Antimicrobials with Polyelectrolyte Nanocomplexes to Treat Bacterial Biofilms and Lung Infections"

**This PDF file includes:**

Figs. S1 to S6

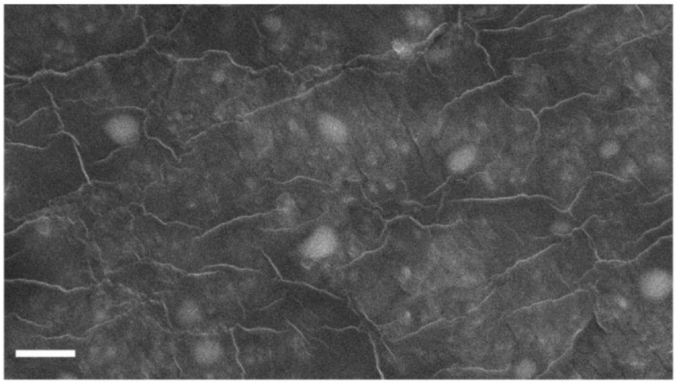

**Fig. S1.**
Scanning electron microscopy image of Tob-NC **1c**. Scale bar = 1 μm

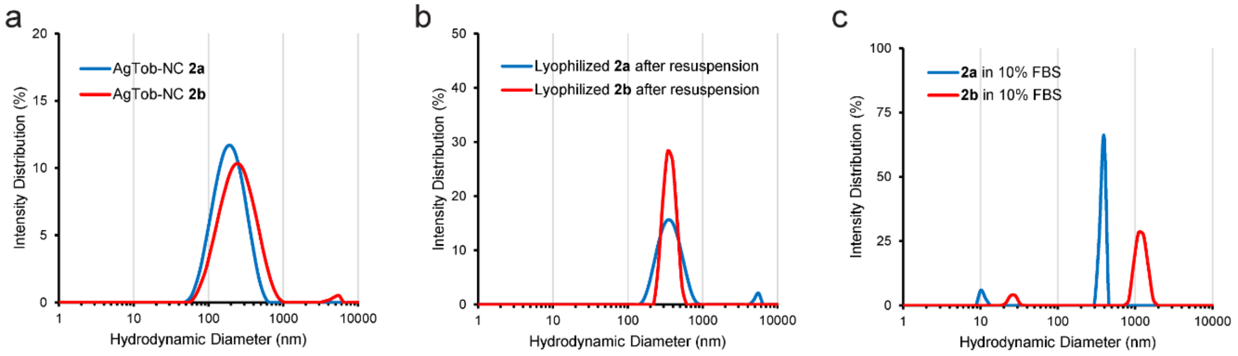


**Fig. S2.**

Storage and stability of AgTob-NCs. Representative intensity distribution curves of AgTob-NCs **2a** and **2b**, (a) immediately after fabrication, (b) after resuspension from lyophilized powder, and (c) after 20 h incubation in 10% FBS solution at 37 °C. Slight increases in diameter were observed after resuspension of lyophilized powder for both **2a** and **2b**. After 20 h in FBS, **2a** size remained stable, while **2b** displayed slight aggregation into ~1 µm diameter particles. In the FBS solutions, small particles were observed at 10-20 nm in diameter, likely corresponding to released AgNP from the NCs. AgNP release was confirmed via detection of UV-Vis absorbance at 390 nm in the supernatant following centrifugation at 13,000 rpm for 5 min.

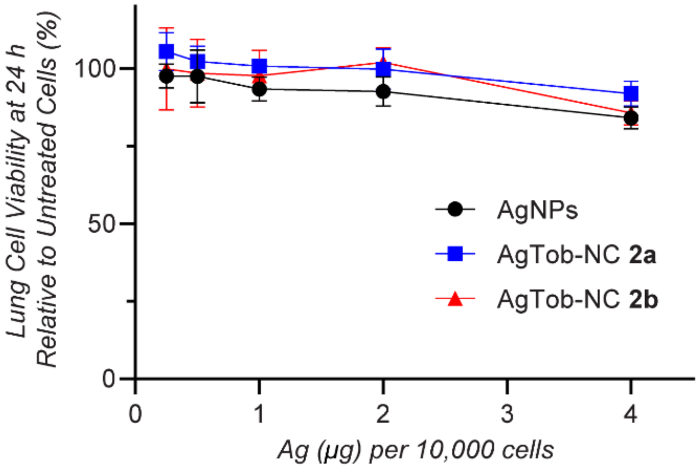


**Fig. S3.**

Biocompatibility studies of AgTob-NCs and AgNP controls. A549 lung cells were exposed to varying doses of AgNPs and NCs for 24 h. Modest decreases in viability were observed at the highest dose tested, corresponding to ~85% viability, as measured by PrestoBlue assays. No statistical significance was observed between groups.

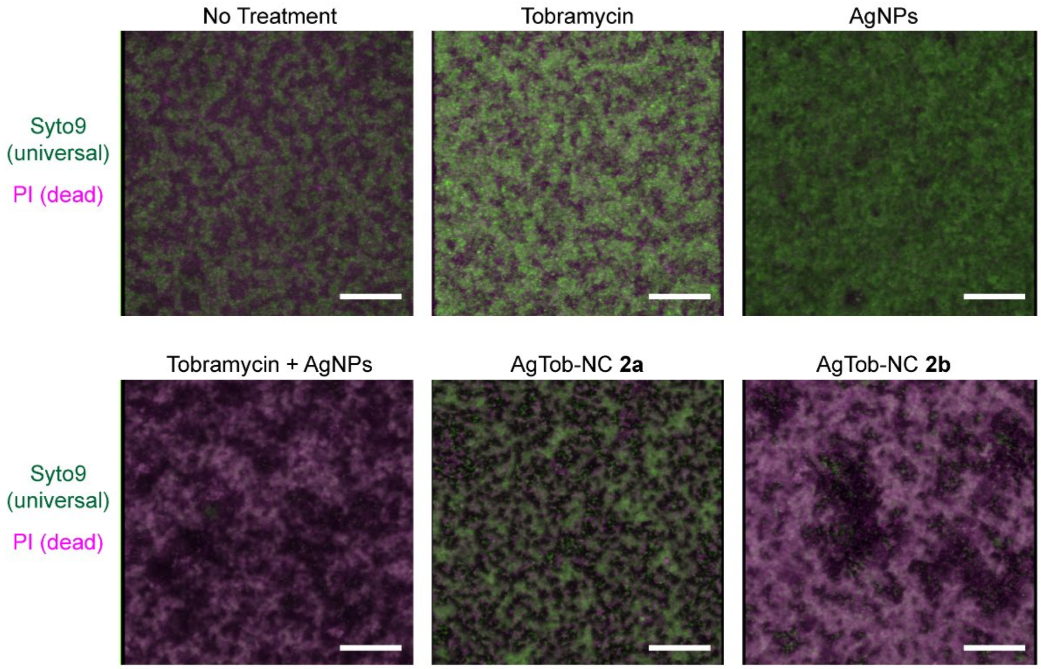


**Fig. S4.**

Antibiofilm activities of AgTob-NCs and controls. Representative 3D projections of confocal fluorescence microscopy images for all groups tested for antibiofilm activity against PA14 biofilms. Syto9 (green) is a universal stain for both live and dead bacteria, while PI (magenta) stains only dead bacteria. White indicates overlay at approximately equal fluorescence intensities. Neither tobramycin nor AgNPs alone demonstrated significant antibiofilm activities, while combination therapeutics in solution or loaded within NC formulations were able to disrupt biofilm biomass, with NC **2b** demonstrating the most pronounced antibiofilm and bactericidal activities. All biofilms were treated with 6.4 µg of Tob and/or 3.2 µg AgNP for 20 h. Images acquired at 20x magnification. Scale bars = 50 µm.

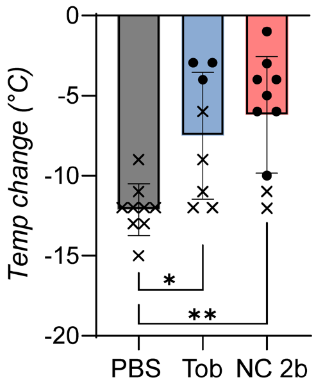


**Fig. S5.**

Body temperature changes over the course of the lung infection survival study. Body temperature was monitored every 2 h over the course of the 24 h study, with final reported temperatures plotted. Temperature data from mice that did not survive are represented as x while data from mice that survived are reported as solid circles. * $p<0.05$, ** $p<0.01$.

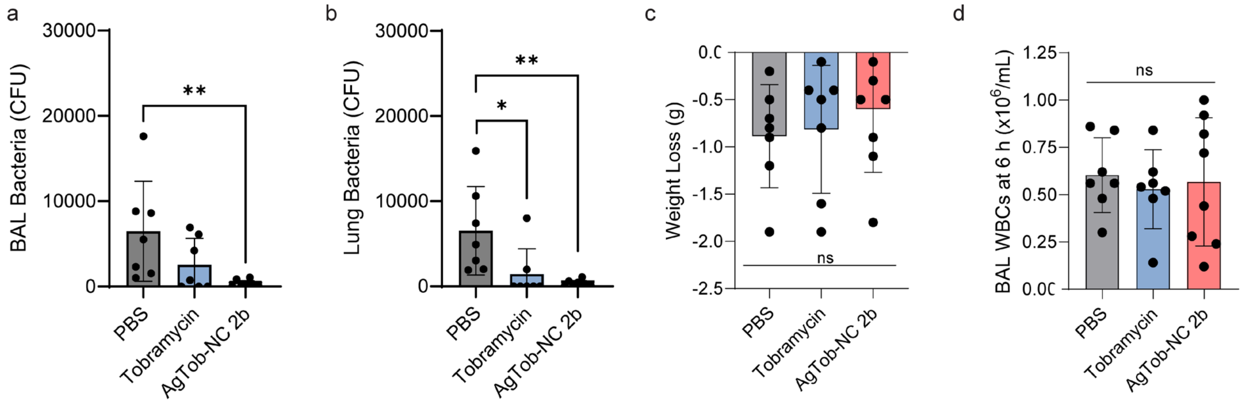


**Fig. S6.**

Bacterial burden and physiological markers of infection at 6 h. Bacterial load in colony forming unit (CFU) for mice at 6 h as determined by (a) bronchoalveolar lavage (BAL) and (b) lung homogenate. (c) Weight loss and (d) white blood cell (WBC) counts revealed no statistical significance between groups at 6 h. * $p<0.05$, ** $p<0.01$.
